# Predictive coding model can detect novelty on different levels of representation hierarchy

**DOI:** 10.1101/2024.06.10.597876

**Authors:** T. Ed Li, Mufeng Tang, Rafal Bogacz

## Abstract

Novelty detection, also known as familiarity discrimination or recognition memory, refers to the ability to distinguish whether a stimulus has been seen before. It has been hypothesized that novelty detection can naturally arise within networks that store memory or learn efficient neural representation, because these networks already store information on familiar stimuli. However, computational models instantiating this hypothesis have not been shown to reproduce high capacity of human recognition memory, so it is unclear if this hypothesis is feasible. This paper demonstrates that predictive coding, which is an established model previously shown to effectively support representation learning and memory, can also naturally discriminate novelty with high capacity. Predictive coding model includes neurons encoding prediction errors, and we show that these neurons produce higher activity for novel stimuli, so that the novelty can be decoded from their activity. Moreover, the hierarchical predictive coding networks uniquely perform novelty detection at varying abstraction levels across the hierarchy, i.e., they can detect both novel low-level features, and novel higher-level objects. Overall, we unify novelty detection, associative memory, and representation learning within a single computational framework.

## 1 Introduction

Humans have an incredible capacity to detect novel stimuli. A classical study shows that human participants can perceive 10000 images and still be able to correctly identify the familiar image in a pair of novel and seen stimuli with 83% accuracy (Standing, 1973). This astounding capacity for *novelty detection (ND)*, also known as familiarity discrimination or recognition memory, is vital for guiding flexible intelligent behavior, such as optimal exploration (Wang et al., 2024).

ND relies on brain regions previously shown to be involved in memory and perception, such as hippocampus, perirhinal and inferotemporal cortex (Brown and Aggleton, 2001). These regions include *repetition suppression* neurons that are most active when presented with novel stimuli, and gradually decline in activity through repeated exposure (Xiang and Brown, 1998; Meyer and Rust, 2018). While their existence is well-documented (Rolls et al., 2004; Suzuki, 1999; Brown and Aggleton, 2001; Viskontas et al., 2006), how these novelty responses arise remains elusive.

Diverse computational models of ND have been proposed. We summarize the two main approaches to ND here, and compare them in more detail in the Discussion. The first approach is developing models specialized just for ND (Bogacz et al., 1999, 2001). One of these models, the Anti-Hebbian model (Bogacz and Brown, 2003a), has been shown to replicate the capacity seen in human recognition memory, when presented with input patterns with a correlation structure likely in neurons representing visual stimuli (Androulidakis et al., 2008; Kazanovich and Borisyuk, 2021; Read et al., 2024). The other approach suggests that ND does not need dedicated circuits, because it can naturally arise within networks that store memory or learn efficient neural representation, as these networks already contain information about the familiar stimuli (Li et al., 1993; Norman and O’Reilly, 2003; Sohal and Hasselmo, 2000). Using existing circuits for ND would reduce brain’s size and energy requirements hence is likely to be favoured by evolution. However, published models combining ND with representation learning do not have high capacity when the input patterns have a biologically realistic correlation structure (Bogacz and Brown, 2003b). Thus in summary, although the hypothesis that ND naturally arises in networks performing other functions is very appealing, the existence of such a combined model that discriminates novelty of correlated patterns with high capacity has not been yet established.

This paper demonstrates that ND naturally arises in predictive coding networks (PCNs), which have previously been shown to effectively learn representations of sensory stimuli (Rao and Ballard, 1999), and support associative memory (AM) (Salvatori et al.,2021). An important feature of PCNs is that they rely on local synaptic plasticity rules, where the weight modification depends only on the activities of pre-synaptic and post-synaptic neurons. PCNs include prediction error neurons that compute the difference between the activity of a particular neuron, and a prediction based on the activity of other neurons. We demonstrate that these error neurons have higher activity for novel stimuli, and it becomes closer to zero as the stimuli are repeated, paralleling the repetition suppression seen in cortical regions underlying ND. We also show that the novelty signal decoded from the prediction error neurons can be used to discriminate the novelty of natural images with a capacity similar to that seen in human experiments. In particular, we show that PCNs are robust to pixel correlation, unlike some of the earlier ND models that perform poorly in images with correlated pixels. To explain this robustness to correlation, we performed a mathematical analysis of a tractable version of PCN, called recurrent PCNs (rPCNs) (Tang et al., 2023b), revealing that rPCNs employ a linear transformation of the covariance structure of inputs that facilitates the discrimination between familiarity and novelty. We also explore hierarchical PCN (hPCN) (Salvatori et al., 2021) in ND tasks and discover that hPCN performs ND for features at varying abstraction levels. Specifically, while the sensory layer of an hPCN can detect the pixel-wise novelty of an image, its hidden layers can detect the novelty of the abstract object in the image by forming latent representations of observed objects.

Overall, ND through predictive coding brings many previous hypotheses about the relationship between ND and other functions like AM (Bussey et al., 2005) and representation learning (Li et al., 1993; Buckley and Gaffan, 1998; Murray and Bussey, 1999; Bussey and Saksida, 2002) to fruition, by providing both a proof-of-concept and a concrete computational framework to test hypotheses on ND and related functions. To our knowledge, no previous computational models have achieved this while maintaining other properties of PCNs such as local learning rules and high capacity. Our models can explain many existing experimental phenomena including the existence of novelty neurons across the visual processing hierarchy and ND’s much larger capacity compared to AM, produce falsifiable predictions for further experiments, and provide a computational framework to ground discussions on the precise relationships between ND, AM, and representation learning.

## 2 Models

In this work, we follow an *energy-based approach* to apply energy-based models for ND tasks. As its name suggests, an energy-based model adjusts its parameters to minimize an energy function when exposed to a pattern, thereby ‘learning’ or ‘memorizing’ it. Then, after training, the energy value of the model evaluated at the query indicates its familiarity — with a lower value signaling familiarity and vice versa, and thus aligning with repetition suppression seen in the brain (Fig. 1). Previous work (Bogacz et al., 1999; Greve et al., 2009) has applied this approach to Hopfield Network (HN) (Hopfield, 1982), a recurrent neural network model for AM, and shows that it successfully performs ND for binary patterns. Here, we extend this energy-based approach to PCNs (Friston, 2005; Tang et al., 2023b) as well as the Modern Continuous Hopfield Network (MCHN) (Ramsauer et al., 2020). Intuitively, energy-based models learn a stimulus by adjusting its weights to minimize the energy (loss) function on that stimulus, which measures ‘surprisal’ (Clark, 2015) of that stimulus to the model. Thus, after training, a familiar stimulus should, on average, have a lower energy or surprisal for the model’s learned weights.

**Figure 1.**
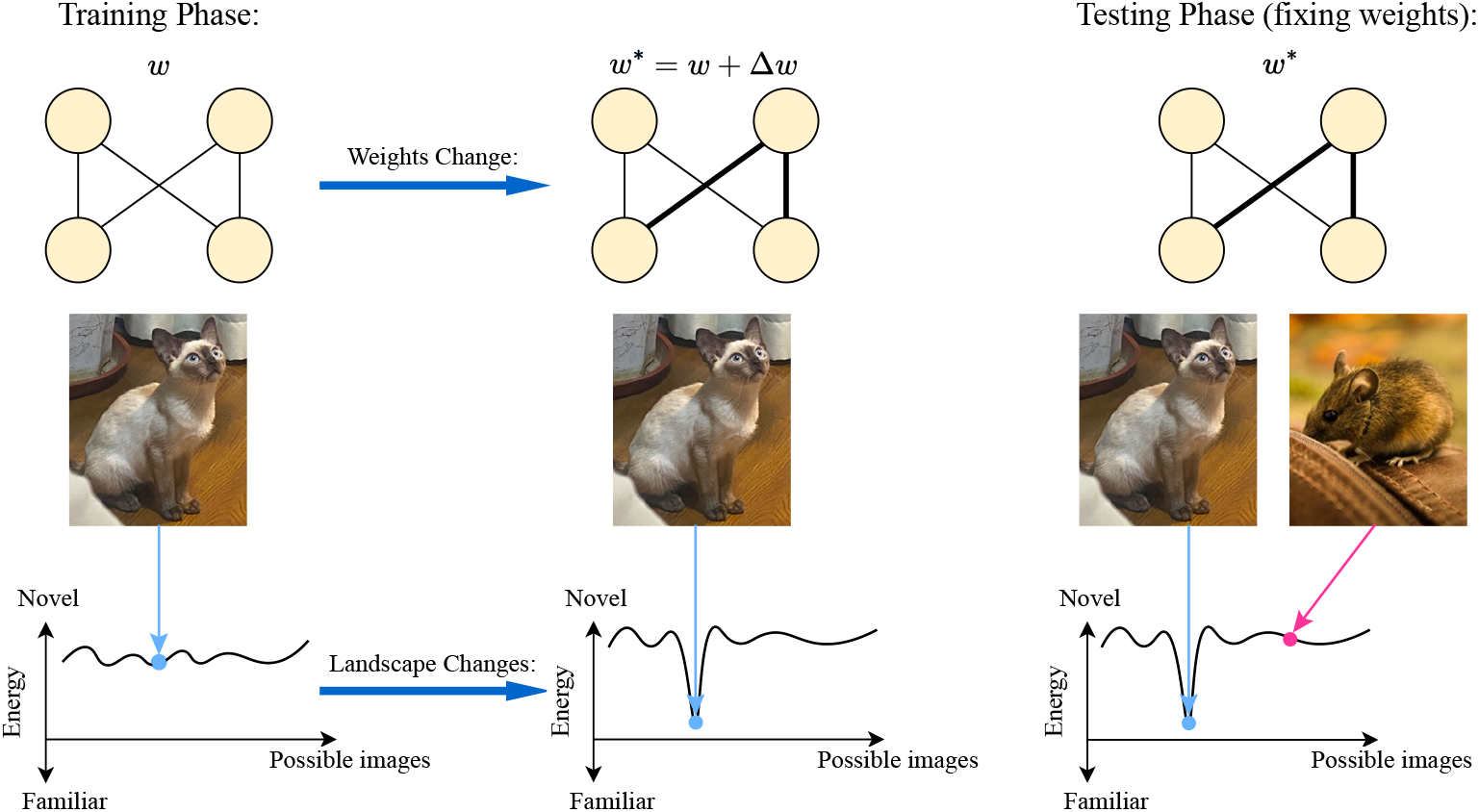
An illustration of the general energy-based approach to ND. During training, an energy-based model (i.e., a model that minimizes an energy function) modifies its weights or parameters to memorize a pattern, which may become a local minimum in the altered energy landscape; this later allows us to simply use the energy value of a pattern as a novelty signal for the memorized pattern or similar patterns in a local neighborhood. The space of possible images is represented in a single dimension on the x-axis for simplicity.

Formally, assume a total of *N*, *d*-dimensional patterns (**x**(1), **x**(2), …, **x**(*N*)) are independent and identically generated from a certain probability distribution. During the *training phase*, patterns to be memorized (which form the columns of the data matrix **X** ∈ ℝ^*d×N*^) are fed to the model one by one, modifying the weights or parameters to reduce the energy. Then, in the *testing phase*, a single *d*-dimensional query pattern **q** is provided to the network, and performing ND only requires reading out the value of the energy function, which indicates the familiarity of the query. This is a general approach that applies to any energy-based model. Even for models that do not use weights explicitly (such as MCHN (Ramsauer et al., 2020)), the energy landscape changes during training as shown in Fig. 1.

### 2.1 Predictive coding networks for ND

The general idea of predictive coding model is that the brain constantly tries to predict the incoming input it receives using the generative model it has learned. The model compares the input with the prediction by calculating their difference in the activity of the corresponding error neurons. The algorithmic goal of PCNs is to minimize the activities of these prediction error neurons (Clark, 2015; Bogacz, 2017) (or dendritic architectures encoding errors (Mikulasch et al., 2023)) by modifying neural activities and synaptic strengths. In this work, we investigate PCNs where the predictions are generated by either recurrent or hierarchical connections.

#### Recurrent predictive coding network

The recurrent PCN (rPCN) is a single-layer neural network model inspired by the recurrent connections of the hippocampus and was originally designed to perform AM tasks (Tang et al., 2023b). We study ND in rPCN in the first part of the Results, because rPCNs are simpler than multi-layer PCNs, and thus analytically tractable.

To illustrate the model, consider a simple 2-dimensional rPCN shown in Fig. 2. When trying to predict the incoming input in value neurons (e.g., level of activity in *x*_1_), rPCN employs lateral connections (*W*) such that its prediction is a sum of a lateral component (e.g., *W*_12_*x*_2_) and a top-down component (e.g., a bias input ***ν***_1_). Then, the corresponding prediction error is *ε*_1_ := *x*_1_ − *W*_12_*x*_2_ − ***ν***_1_, and can be computed by error neurons receiving the connections shown in Fig. 2 (connections from error *ε*_*i*_ to value neurons *x*_*i*_ are not used in ND but are included in the figure as they are useful for memory retrieval rPCN can perform (Tang et al., 2023b)). To reduce these errors during learning, the model modifies its synaptic weights *W* and ***ν*** to reduce the squared prediction errors. During the *training phase*, rPCN’s dynamics aim to minimize the total squared prediction errors *ε*_*i*_ of all neurons with respect to *W* and ***ν***. This algorithmic goal corresponds to the minimization of the following energy function:

**Figure 2.**
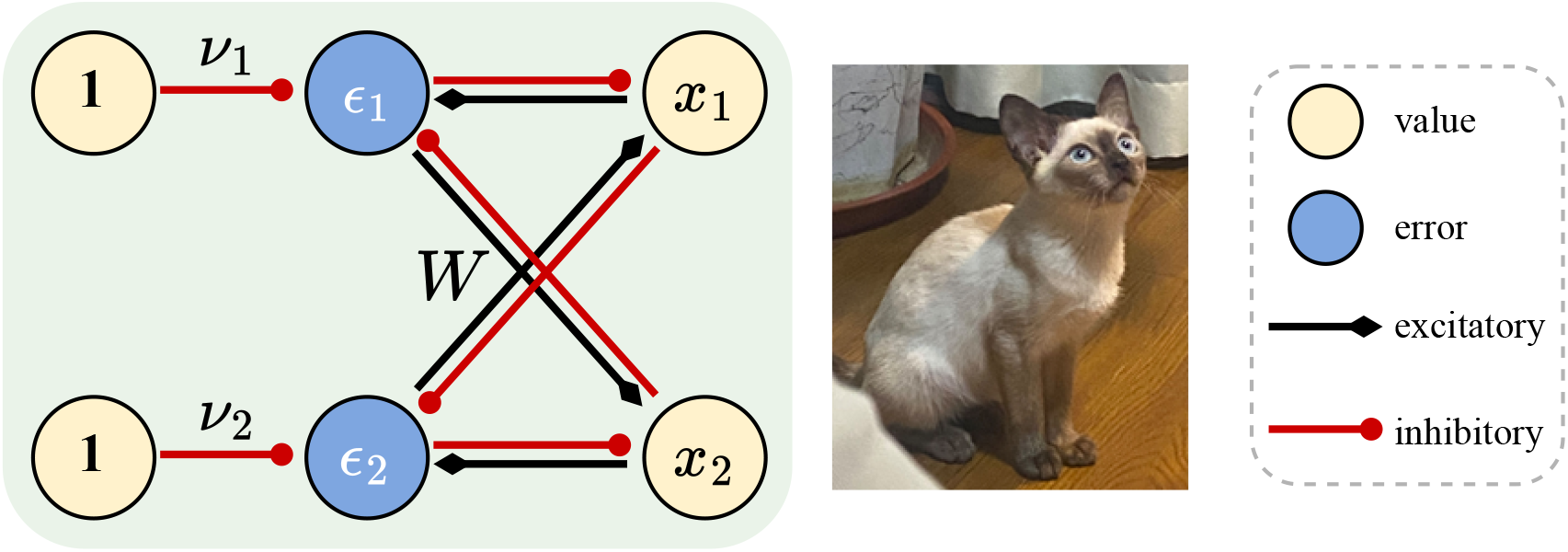
Recurrent predictive coding network (rPCN). The illustration is simplified to *d* = 2 dimensional. At training, the **x** = (*x*_1_, …, *x*_*d*_) is fixed to the values of the image while all other values and parameters change according to the learning rules.

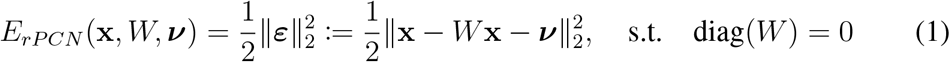

where *W* is the weight matrix implicitly encoding covariance and ***ν*** is the bias vector (Tang et al., 2023b). The optimization is subject to the constraint that *W* has zero diagonal to prevent the trivial solution *W* = *I* (corresponding to the absence of self-connections in Fig. 2). The zero diagonal will stay zero throughout training. rPCN updates its parameters *W* and ***ν*** with gradient descent by calculating the gradient of *E*_*rPCN*_ based on the training images 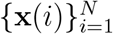:

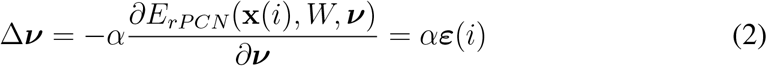

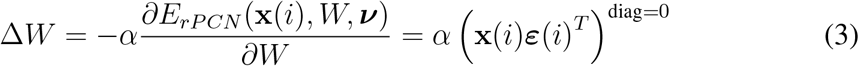

where ***ε***(*i*) := **x**(*i*) − *W* **x**(*i*) − ***ν*** is the prediction error of the *i*-th training image and (·)^diag=0^ denotes that the diagonal elements are enforced to be 0, maintaining the unchanged zero diagonal elements of *W*, and *α* is the learning rate parameter. In numerical simulations, rPCN can also be trained efficiently using the batch version of these learning rules. At the *testing phase* for a query image **q**, rPCN initializes the activity to **q**; then, it evaluates the energy function on **q** as the novelty signal:

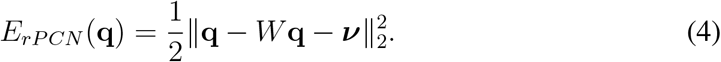

Notice that the weight update rules are Hebbian and require only local computations. For instance, since Δ***ν***_1_ = *αε*_1_ and Δ*W*_12_ = *αx*_2_*ε*_1_, the learning rules for ***ν*** and *W* are both a product of their respective pre- and post-synaptic activities (Fig. 2). The energy can be evaluated by summing up activities of error neurons, following mechanisms such as those discussed in Carandini et al. (2005), or simply by an external algorithm decoding the population activity of neuron assemblies. rPCN is also more numerically stable than other formulations of PCNs with recurrent connections (Tang et al., 2023b).

#### Hierarchical predictive coding network

Most existing models for ND have been proposed to account for the detection of novel individual elements of provided inputs. For example, if a model is trained on an image of a Siamese cat, an image with only a few pixel values changed from the original image will be detected as a novelty by classical ND models, although the semantic meaning i.e., a Siamese cat, is not changed by altering these few pixels. By contrast, the more flexible detection of semantic or abstract novelty is closer to the one that animals employ effortlessly daily to guide behaviors like exploration.

Hierarchical predictive coding is natural for this purpose as it detects features of increasing abstraction levels across the hierarchy, mirroring the role performed by the ventral visual stream and the visual cortex (Rao and Ballard, 1999; Bussey and Saksida, 2002). Therefore, to fill in this gap we investigate ND in hierarchical PCN (hPCN) (Salvatori et al., 2021) illustrated in Fig. 3. In hPCN, neurons in layer *l* are denoted as vector **x**^*l*^ and its value is compared against top-down prediction from the neurons in *l* + 1 transformed by weights *W*^*l*+1^ to produce a corresponding error signal ***ε***^*l*^ defined as

**Figure 3.**
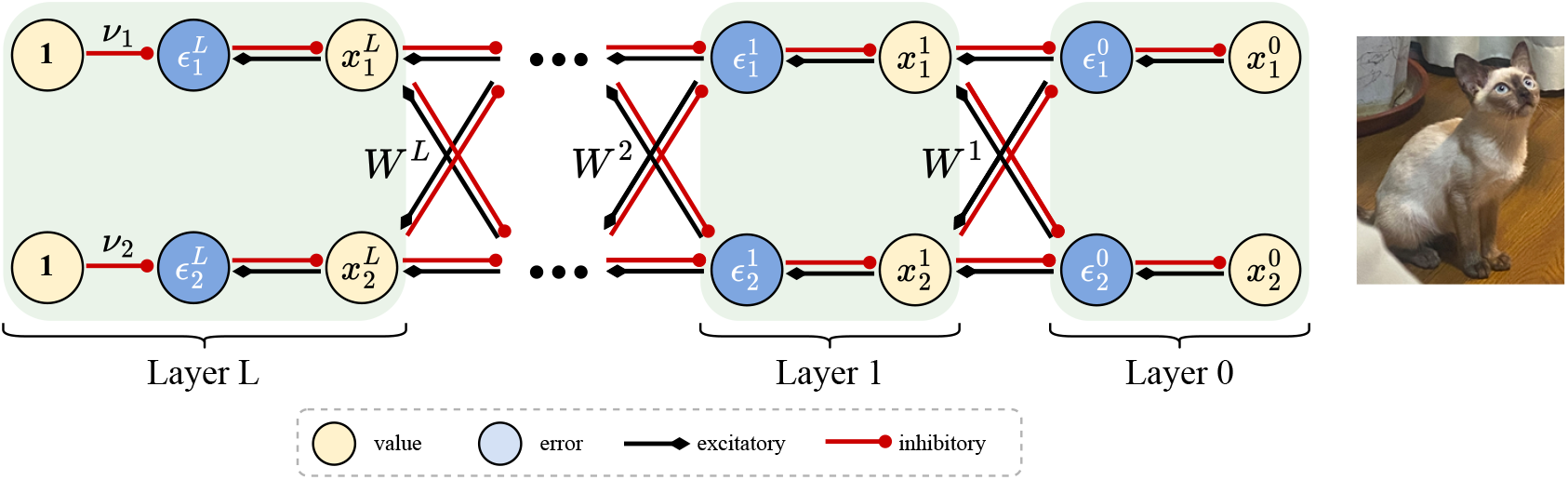
Hierarchical PCN (hPCN). A 2-dimensional, *L*-layer hPCN model from Salvatori et al. (2021). Layer 0 is the sensory layer where input patterns enter the model.

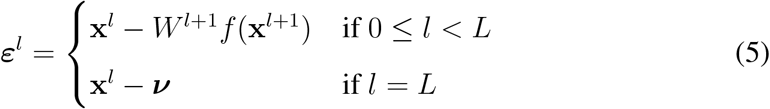

where *f* is an element-wise nonlinear function. The optimization goal of hPCN is then to minimize the sum of squares of all energy units:

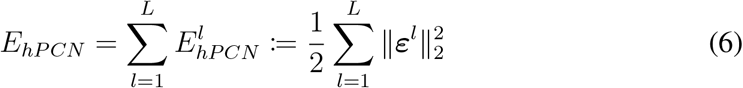

where 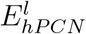 denotes the energy (or half of the sum of squared errors) at layer *l*. During training, **x**^0^ is fixed at the input to memorize, and the activity of value neurons in hPCN is modified until convergence according to:

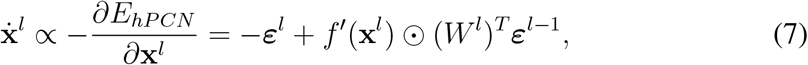

where ⊙ denotes the element-wise product, and *f* ^*′*^ the derivative of the nonlinear function. When the activities converge, the weights are then modified:

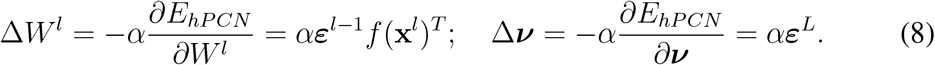

To use hPCN for ND, **x**^0^ is fixed to the query **q**, and the model performs inference following Eq. 7 again until convergence. Then, the layer-wise energy values 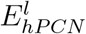 (instead of the total energy *E*_*hPCN*_) at all layers *l* = 0, …, *L* are evaluated. The key idea here is that since 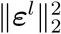 serves as a novelty signal for the feature detected by value neuron **x**^*l*^, 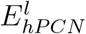 can serve as a novelty signal for features collectively learned by layer *l*. Therefore, different layers can signal different levels of novelty by learning features of different abstraction levels and encoding their novelty by layer-wise energy functions. Importantly, learning (Eq. 8) in hPCNs is also Hebbian, and inference dynamics in the model only require local information. In this work, we also imposed local connectivity constraints between early layers of our hPCN model to mimic the limited receptive fields that neurons in the early processing hierarchy tend to have (details in the Results section).

### 2.2 Hopfield networks for novelty detection

To put the results of simulations of PCNs into context we compare their capacity for ND with HNs. HN is a classical energy-based model for AM (Hopfield, 1982), which has also been shown to demonstrate the capacity of ND by measuring the degree of familiarity via its energy function after training (Bogacz et al., 1999). It has an energy function

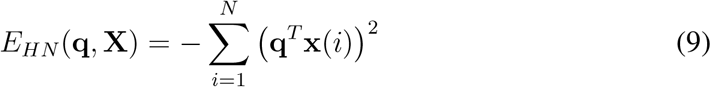

This energy can also be rewritten in terms of covariance matrix of patterns ∑:

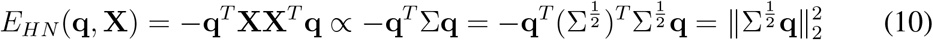

In this work, we extend this approach to MCHN (Ramsauer et al., 2020), a modification of the original HN that performs AM effectively for images with continuous (rather than binary) pixel values. MCHN has an energy function

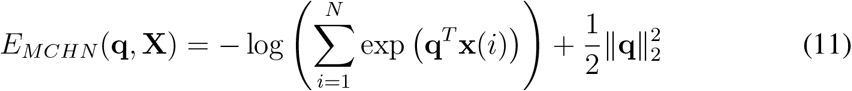

Past research has proposed biologically plausible implementations for HN, MCHN, and even more generally Universal Hopfield Network (UHN) (Millidge et al., 2022). Please note that, the energy value in Eqs. 9 and 11 are both functions of the training set (**X**). They differ from rPCN (Eq. 4) and hPCN (Eqs. 5 and 6), whose energy is a function of the model weights after training (*W*). As such, Eqs. 9 and 11 allowing us to skip training HNs and to evaluate the energy (novelty signal) directly from the training set (given query **q**). Thus, it is important to note that under this experiment setup, the performances of HN and MCHN are given a tiny advantage over PCNs as they do not require gradient-based training of any weights, which may introduce noise to PCNs.

## 3 Results

### 3.1 Predictive coding replicates repetition suppression

We first investigate whether the activities of error neurons in PCNs will present repetition suppression upon multiple exposures to the same stimulus. We trained both an rPCN and an hPCN with 2 hidden layers of 256 neurons on one single image extracted from the Tiny ImageNet (Deng et al., 2009) dataset and recorded the mean of squared error neuron activities as well as the distribution of the absolute activities across the error neuron populations. The results are shown in Fig 4. It is not surprising that the overall activities of error neurons reproduce repetition suppression, as PCNs are iteratively trained to minimize prediction errors. Importantly, however, this result suggests a possible mechanism underlying repetition suppression signaling novelty in the cortex, which stems from the minimization of local prediction errors or energies. It is also worth mentioning that we do not constrain the error neurons to be non-negative in our models. However, it has been suggested that positive and negative prediction errors are encoded by separate groups of neurons in the cortex, signaling novel stimuli that are ‘stronger’ or ‘weaker’ than predicted (Keller and Mrsic-Flogel, 2018). In such an architecture, the energy can be simply read out as a sum of non-linearly transformed (squared) activity of the prediction error neurons.

**Figure 4.**
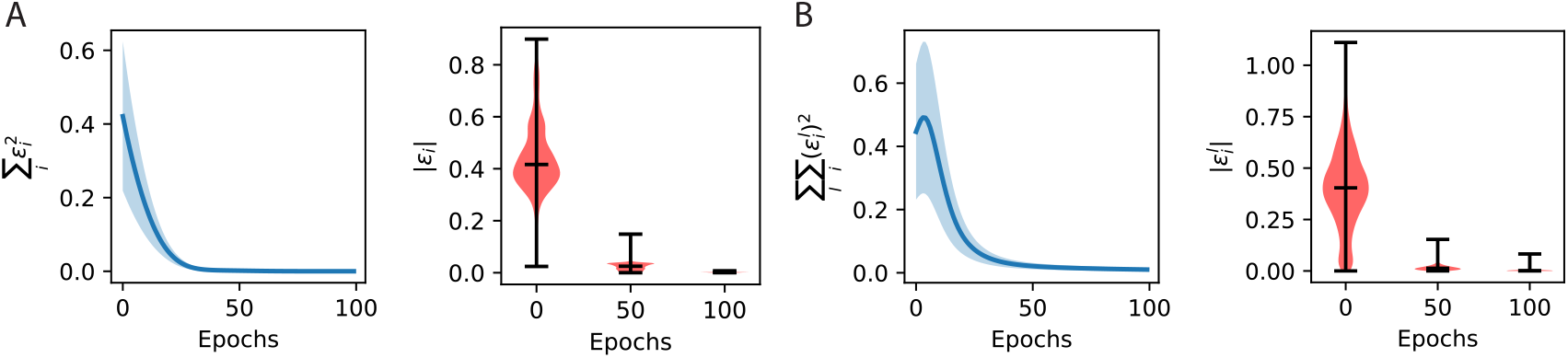
Activites of error neurons throughout training. A: Evolution of total energy (left) and distribution of absolute error neuron activities throughout the training of an rPCN on one image from Tiny ImageNet, with a learning rate of *α* = 0.0003. B: Same as A, but for an hPCN with 2 hidden layers, with a learning rate of *α* = 0.0003.

### 3.2 Predictive coding replicates human novelty detection abilities

Here, we demonstrate that rPCNs can discriminate familiarity for a large number of stimuli and can match the experimentally observed capacity of human recognition memory (Standing, 1973). We examine ND in rPCN and control models (HN, and MCHN) using three different datasets: 500-dimensional Gaussian images with uncorrelated and correlated pixels (with a 0.4 covariance between any two pixels), and 64 × 64 images from the Tiny ImageNet dataset (Deng et al., 2009). We say a dataset has correlated pixels if its feature-by-feature covariance matrix is non-diagonal. We include pixel correlation as a benchmark since past works have found that correlated pixels often pose challenges to ND algorithms (Bogacz and Brown, 2003b). Moreover, such robustness is important since natural images have highly correlated pixel values (e.g., if a pixel is dark, then pixels close to it tend to be dark as well). Additionally, the activity of neurons representing stimuli in higher visual areas (e.g., the perirhinal cortex) was also observed to be correlated between trials on which different stimuli were presented (Erickson et al., 2000). For rPCN, the model is first trained on a certain number of patterns or images following Eqs 2 and 3, and the energy function (Eq 4) is then calculated, given a novel or familiar **q**, to evaluate the model’s detection of novelty. For HNs, since effectively there is no ‘training’ phase of the model, Eqs 9 and 11 are directly used to evaluate the performance (detailed experimental procedures, including the calculation of error probabilities, are provided in Appendix B).

The performances are shown in Fig. 5, where we also compare all three models with the experimentally observed performance of humans in discriminating familiarity of natural images (Standing, 1973). The first row displays the average error probability as a function of the number of presented patterns or stimuli and the second row shows the number of patterns retained in memory (defined in Standing (1973) and given in Appendix B). With both axes logarithmic, the plots reveal a power-law relationship observed for human participants (Standing, 1973). In the leftmost column, pixels are uncorrelated and the three models have similar, decent accuracy. Importantly, when features become correlated in the middle column, the rPCN keeps a similar level of performance. By contrast, the error of HN and MCHN greatly increases, approaching chance-level performance for a larger number of patterns, as seen for HN in a previous study (Androulidakis et al., 2008). We then investigated if the observed effect of pattern correlation generalizes to real-world images, which tend to have correlated pixels, especially among pixels that are spatially proximate to each other. This is indeed what testing on the Tiny ImageNet reveals in the right column of Fig. 5: HN and MCHN perform poorly even for a small number of stored patterns. Please note that the performances of HN and MCHN are poor despite being given an advantage in performance, which comes from directly evaluating the energy functions without gradient-based training that is necessary for rPCNs. By contrast, rPCN achieves ND accuracy matching those of human participants (Standing, 1973).

**Figure 5.**
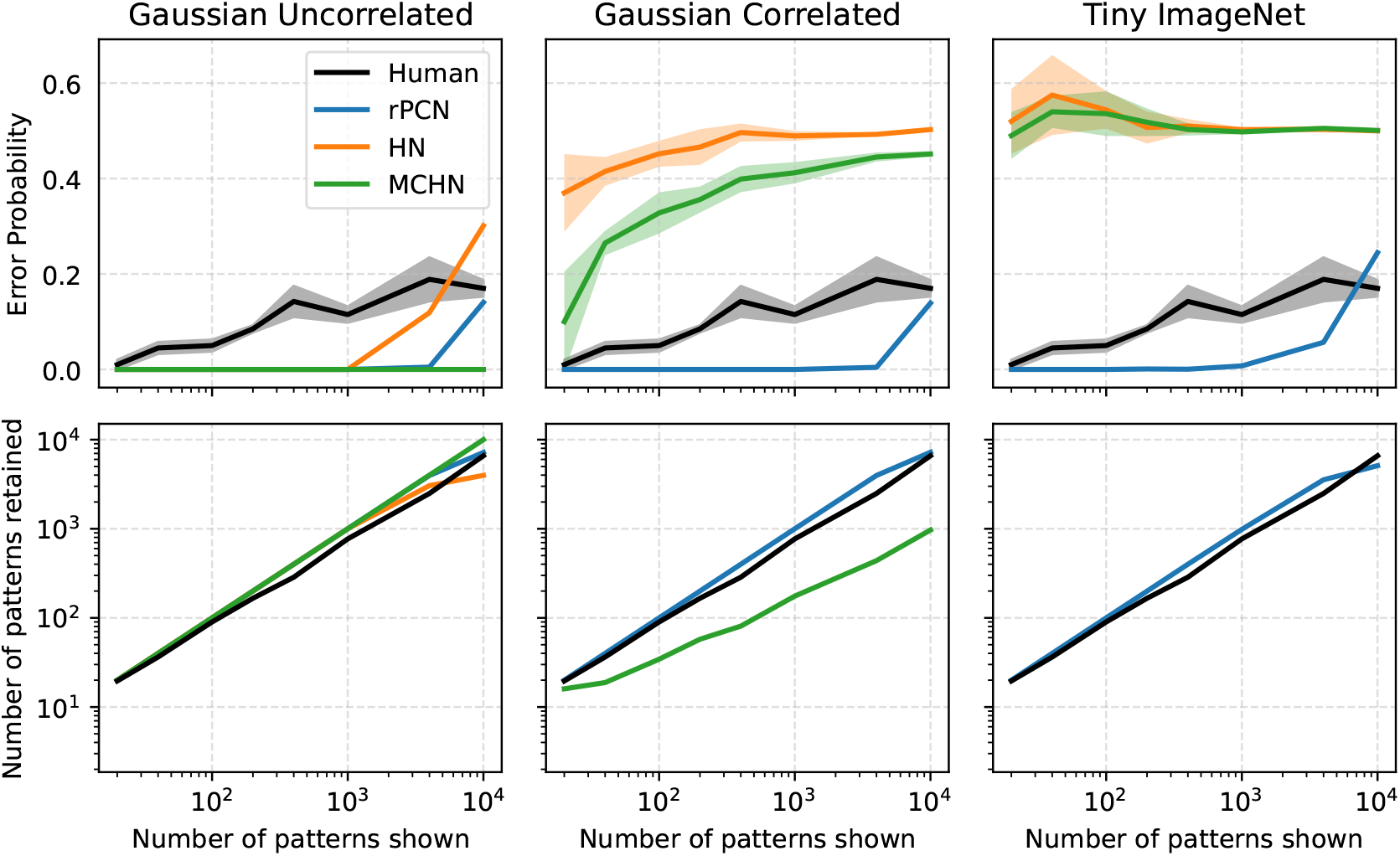
Comparing the performances of rPCN, HN, and MCHN on various datasets. Top row: Error probability as a function of the number of training samples. An error probability of 0.5 corresponds to a baseline level equivalent to guessing. The first two columns are produced with *d* = 500. For the last column, *d* = 4096 for (grayscale) Tiny ImageNet (Deng et al., 2009). All error bars are obtained over 5 trials, matching the number of human participants in the study by Standing (1973). Bottom row: Number of patterns retained in memory as a function of the number of training samples. HN is not shown in the middle plot and both HN and MCHN are not shown in the last plot due to certain data points falling outside of the log-log scale and their performances being close to the chance level.

### 3.3 Novelty as a distance in an embedded space

To provide an understanding for why rPCNs can effectively detect novelty for correlated patterns, while HNs cannot, we analyze mathematically ND in these models. We provide a unifying mathematical account for HN and rPCN in ND, showing that they can both be considered as measuring the Euclidean distance between a query and the mean of the training data, but on different transformed planes. Formally, when the stored patterns **X** have mean 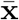, there exists a parameterized class of distance of the form

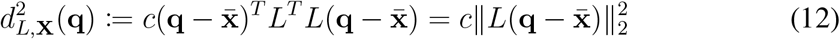

which can be seen as the squared Euclidean distance from the **q** to 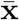 in the space transformed by a *d* × *d* matrix *L* and multiplied by a scalar constant *c*. Using HN for ND can be seen as metric learning of this form:

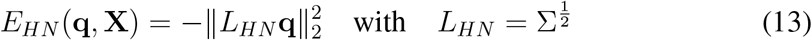

where patterns are assumed to be centered around **0** and ∑ is the covariance matrix of patterns (derivation in section Models). On the other hand, for rPCN the following theorem holds:

**Theorem 1** (rPCN performs metric learning for ND). *When the learning of rPCN has converged and a query* **q** *is supplied at the testing phase, we have*

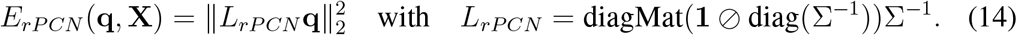

Here, ∑ is the covariance matrix of the memorized patterns, ⊘ is the Hadamard (element-wise) division, **1** denotes the 1-vector, diag extracts the diagonal elements of a matrix and converts them into a *d*-dimensional vector, and diagMat converts a vector into a diagonal matrix. The proof uses Lagrange multipliers and can be found in Appendix A. Combined, Eq. 13 and 14 show that both HN and rPCN can be seen as learning a metric from data through the covariance matrix ∑.

We then investigate the exact transformation *L*_*HN*_ and *L*_*rPCN*_ perform on the distance, by visualizing a simple two-dimensional example. Fig. 6 visualizes the effect of metric learning that HN and rPCN perform specifically when trained on two-dimensional patterns (grey dots). The patterns are randomly drawn from a normal distribution with a positive correlation. The correlation makes ND challenging since a typical familiar point (purple) and a typical novel point (orange) may be equally away from the mean (**0** here, black dot in Fig. 6A) by Euclidean distance. Fig. 6B illustrates that HN first transforms all points from Fig. 6A by 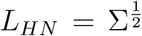, and then measures the *negative* distance to the mean **0** (note the inverted contour color scale). Because of the negative sign, the further the distance (away from the mean), the more familiar the query. Although this appears to address the particular problem of indistinguishable purple and orange dots, it will classify the closest dots to the origin as the most novel. Fig. 6C shows that rPCN also transforms all points from Fig. 6A, but by a different matrix. There, without the negative sign, the further the distance (away from the mean), the more novel the query. Importantly, this alters the covariance structure of the data cloud (i.e., pulling the orange dot from the origin), making it easy to measure familiarity by the distance between the query and the origin.

**Figure 6.**
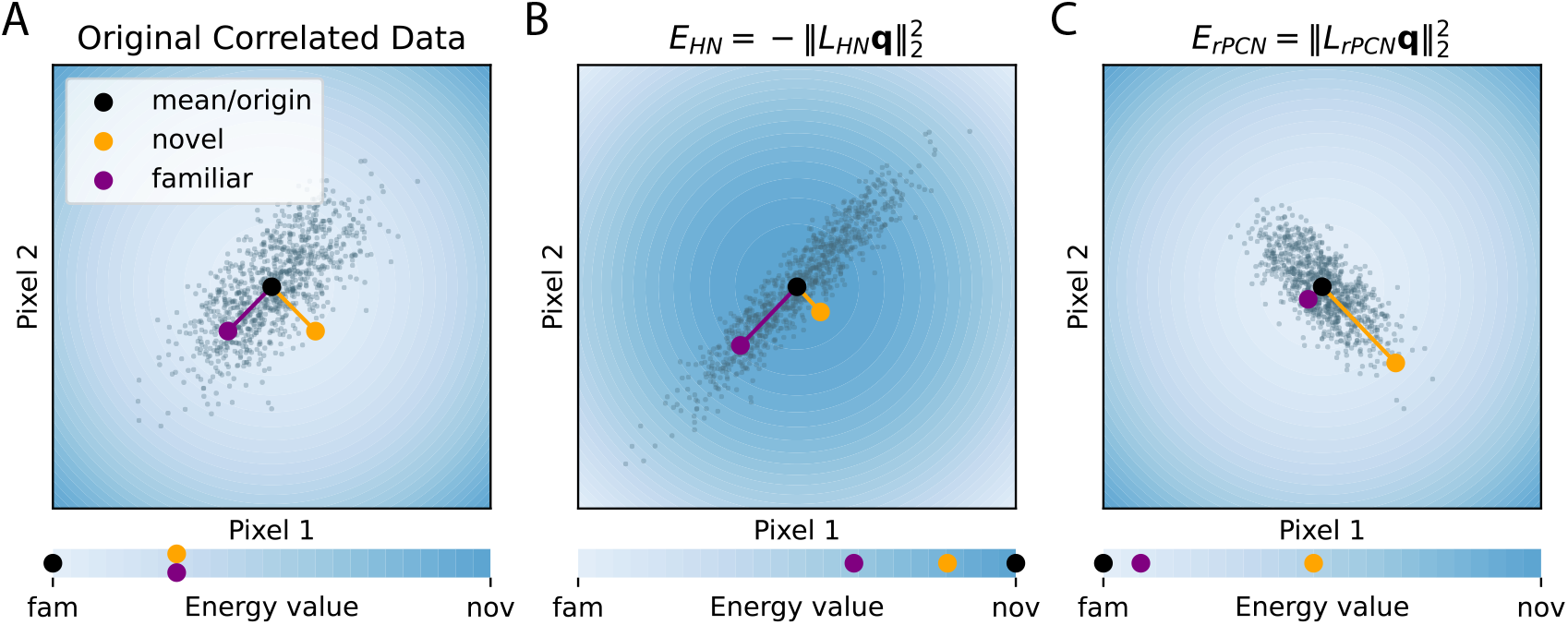
Visualizing the effects of different ND models on 2-dimensional Gaussian data as different linear transformations. The data cloud is generated from a correlated Gaussian distribution where the two pixel values have a covariance of 0.7. In each panel, the space is colored according to the energy value shown in the corresponding bar at the bottom. The energy value of a query pattern **q** in panel A is squared 2-norm (i.e., 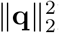), while the energy functions of the corresponding models (HN and rPCN) are used in panels B and C. The energy function of each panel can be seen as a transformed squared 2-norm, each by a different transformation (A: identity matrix *I*_2_; B: *L*_*HN*_ from Eq. 13; C: *L*_*rPCN*_ from Eq. 14). Note that the direction of novelty is inverted in panel B because of the negative sign in Eq. 13.

### 3.4 Multilevel novelty detection

In this section, we distinguish detecting *sensory novelty* from detecting *semantic novelty*. The former is the type of ND that has been addressed so far in this paper—depending on past occurrences, labeling entire images as novel or familiar accordingly based on the individual pixel values. It is also the ND that the overwhelming majority of ND literature focuses on. On the other hand, semantic novelty involves the extraction of abstract features such as semantic or categorical information, such as the numerical digit in an image.

To illustrate their differences, consider the simplified training regimes shown in Fig. 7, where a subject or model is trained on one particular image of the digit ‘4’ only as shown on the left. For sensory (pixel) novelty, the same image of ‘4’ should have a low novelty, but a different image of ‘4’ has a higher novelty value due to its slightly different pixel composition. An image of ‘5’ thus has an even higher value of novelty as its pixel composition deviates further from the image of ‘4’ used in the training. On the other hand, for semantic novelty, both images of ‘4’ have a low novelty value since they both share the same (semantic) feature of ‘4’, with the image of ‘5’ having a high novelty value as before. For animals, the ability to detect novelty for various semantic features is arguably even more important, thus a biologically plausible computational mechanism for such capacity is of great interest to neuroscience.

**Figure 7.**
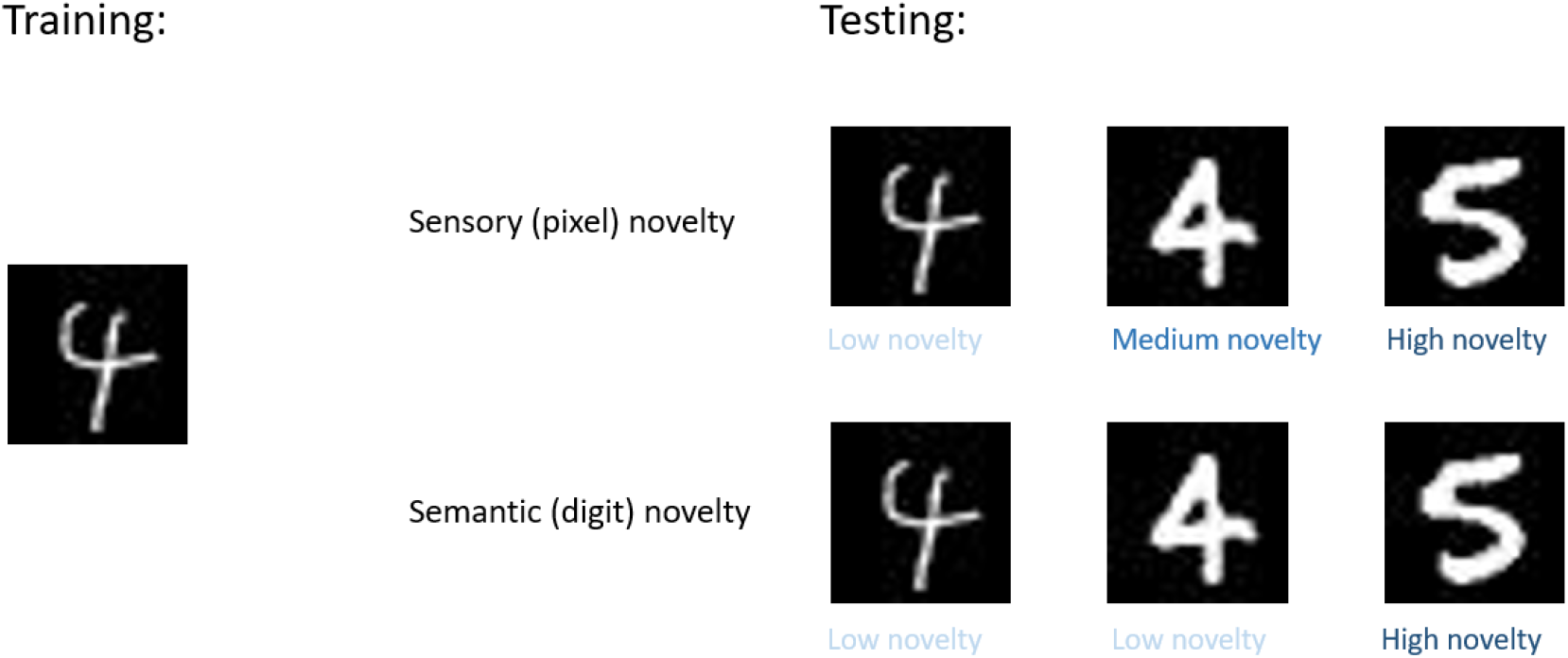
Comparison of sensory and semantic novelty detection in a simplified training regime.

To show how hPCN provides a potential solution, recall that, similar to rPCN, the overall energy function of hPCN in Eq. 5 will be minimized for patterns in the training set (familiar patterns). However, in hPCN, an error neuron on a particular layer *l*, say 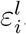, will signal layer-wise novelty of the features represented by the corresponding value neuron 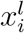 at this layer. For example, at the sensory layer, the error neurons detect ‘How novel is the query at pixel-level’ whereas at higher layers, the error neurons can detect ‘How novel is the (abstract) feature of the query.’ Thus, using layer-wise, rather than the overall energy function can potentially help the detection of novel abstract features.

To test this, we trained a 3-layer hPCN on *N* = 100 images of a particular digit ‘4’. Fig. 8A demonstrates the architecture of this model. Layer 0 is the sensory layer where input patterns enter the model (i.e., fixed to image value during training and testing), with each neuron corresponding to a unique pixel. Instead of letting each neuron in layer 0 be fully connected to every neuron in layer 1 as in Fig. 3, we restrict each (value) neuron in layer 1 to be connected with only a 9 × 9 subset of (error) neurons in layer 0 to mimic the anatomy in early visual areas. We keep layer 2 (value) neurons fully connected to layer 1 (error) neurons. Therefore, a layer 1 neuron has a limited receptive field in a subset of pixels while a layer 2 neuron has the entire latent representation in its receptive field. The model is then trained following the procedure described in the Models section. After training, we test the trained model on three separate sets of queries: (1) the images of ‘4’ that the model was trained on, or ‘familiar ‘4’s’; (2) *N* unseen images of ‘4’, or ‘novel ‘4’s’; (3) *N* (unseen) images of a different ‘5’, or ‘novel ‘5’s’, by fixing layer 0 to these test images and performing inference to minimize the overall energy function in the model until convergence (Eq. 7). Fig. 8B shows the converged distributions (across *N* samples) of energy values 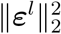 in all 3 layers given the 3 different sets of queries. It can be observed that in all layers the novel ‘5’s are represented with high energy values, whereas the energy difference between familiar and novel ‘4’s decreases as the layer number increases and the energy distributions become similar at the topmost layer. We then quantitatively confirm this trend in Fig. 8C, where we measure the *d*^*′*^ separability, a standard measurement of the separability between two empirical distributions (Grant et al., 2016), between the layer-wise energy distributions given the three sets of stimuli. The *d*^*′*^ value between the energy distributions of familiar and novel ‘4’s monotonically decreases as a function of layer number, while the *d*^*′*^ value between the energy distributions of familiar ‘4’s and novel ‘5’s is consistently higher, suggesting that hPCN is able to detect solely semantic novelty higher in its hierarchy.

**Figure 8.**
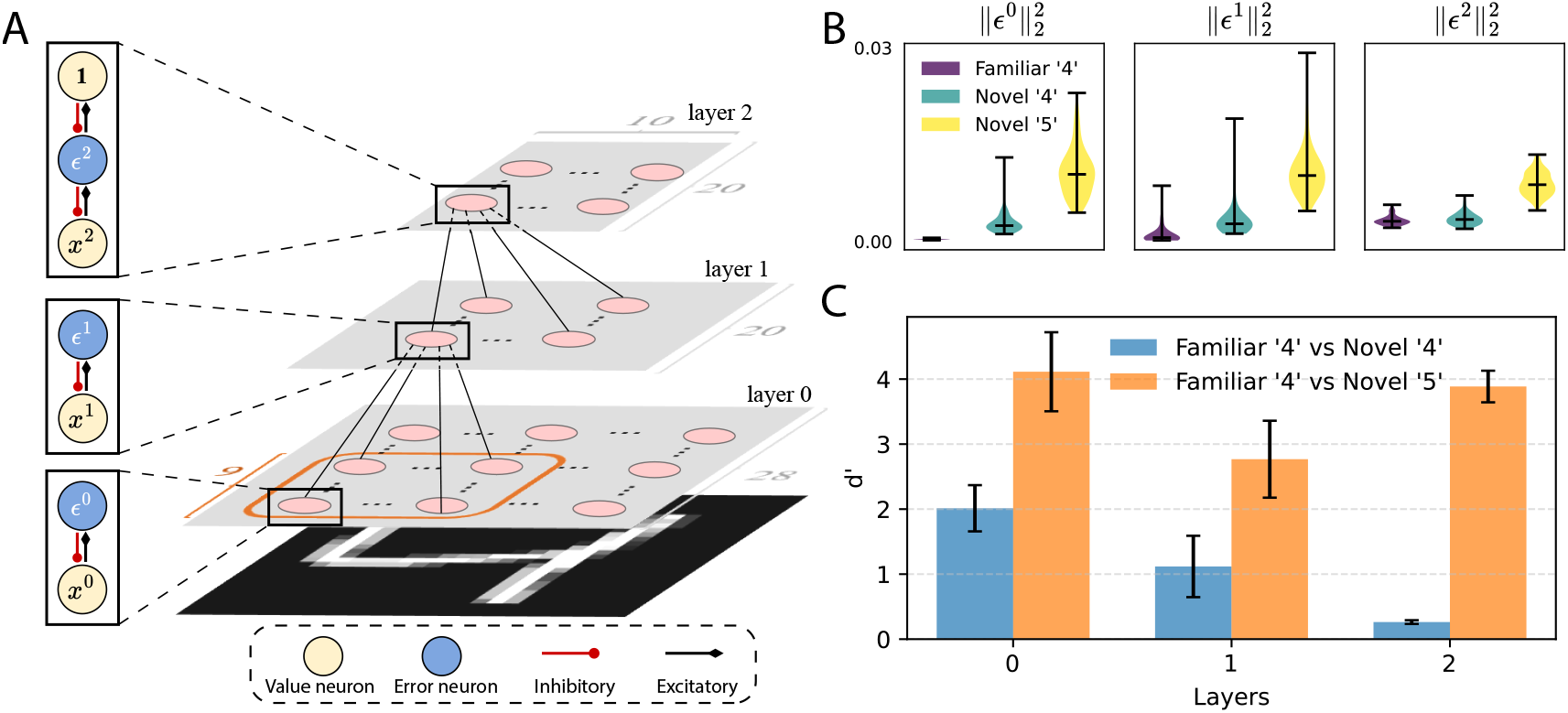
Detecting novelty for features at varying abstraction levels using different layers of error neurons in hPCN. A: Illustration of hPCN with locally connected weights between layers 0 and 1. B. Violin plots of the distribution of layer-wise energy values 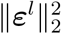 given the three different query sets of size *N* = 100. C: *d*^*′*^ separability score between the empirical distributions in panel B. Error bars obtained with 5 random seeds.

Overall, these results suggest that hPCN develops novelty detection for features of various levels of abstraction. This generalized notion of ND more closely models the ND that animals employ to guide flexible and intelligent behaviors (Bussey and Saksida, 2002).

## 4 Discussion

### 4.1 Relationship to other models of novelty detection

Table 1 compares various ND models in the literature with respect to multiple desired criteria. As mentioned in the Introduction, the first approach to ND is designing specialized models for this task. One example of this approach is the *anti-Hebbian model* (Bogacz and Brown, 2003a), which employs anti-Hebbian learning that weakens connections between layers in response to repeated exposure to the same stimuli (so it uses local learning rules). This model achieves high capacity even when patterns are correlated (Androulidakis et al., 2008). Recently, Kazanovich and Borisyuk (2021) and Read et al. (2024) have extended the anti-Hebbian model and bridged the gap between testing ND on binary patterns (i.e., each pixel value can be either 0 or 1) and natural images. In their experiments, the input to their anti-Hebbian ND model is not the image itself (as is the case in all of our experiments), but rather the features processed and detected by a deep convolutional network. With this model, Kazanovich and Borisyuk (2021) replicated the experimental observation that human subjects perform better for ND tasks on natural images compared to procedurally generated abstract images (Bellhouse-King and Standing, 2007). Another interesting work (Tyulmankov et al., 2022) demonstrated that if meta-parameters of learning rules are trained to optimize ND, the resulting learning rules correspond to those in the anti-Hebbian model, providing another indication of its efficiency. However, anti-Hebbian models have the limitation of being dedicated just to ND so they do not contribute to representation learning and AM.

**Table 1:** Comparison of various ND models across individual criteria. ‘High ND capacity’ is defined as being able to achieve an approximately linear relationship in its number of patterns retained on log-scale for up to 10000 images, generated from an uncorrelated multivariate Gaussian distribution (see bottom left panel on Fig. 5).

|  | Anti-Hebbian | HN & MCHN | Norman & O’Reilly<br>Sohal & Hasslemo | Infomax | Cowell et al. | PCN (ours) |
| --- | --- | --- | --- | --- | --- | --- |
| High ND capacity | ✓ | ✓ | ✓ | ✓ |  | ✓ |
| Robustness for correlated pixels | ✓ |  |  | ✓ |  | ✓ |
| Local learning rules | ✓ | ✓ | ✓ |  |  | ✓ |
| Performs AM |  | ✓ |  |  |  | ✓ |
| Performs representation learning |  |  | ✓ | ✓ |  | ✓ |
| ND at different abstraction levels |  |  |  |  | ✓ | ✓ |

The second approach mentioned in the introduction is designing models that combine ND with other functions. They include HN, as well models combining ND with learning representation (Norman and O’Reilly, 2003; Sohal and Hasselmo, 2000). Particularly, recognizing the close relationships between the hippocampus and neocortex, the neural network model developed by Norman and O’Reilly (2003) for ND aims to disentangle the hippocampal and neocortical contributions. In comparison, Sohal and Hasselmo (2000) more specifically targets the repetition suppression behaviors in the inferotemporal cortex. However, these combined models do not have high capacity when input patterns are correlated (Bogacz and Brown, 2003b). To illustrate that it is theoretically possible to effectively detect novelty in a network that learns representation, Lulham et al. (2011) showed that neural networks implementing the Infomax algorithm (Bell and Sejnowski, 1995) have a large capacity for ND and robustness to correlated inputs. However, these networks are trained with non-local learning rules, which greatly limits their biological plausibility.

A different approach to modeling recognition memory was taken by Cowell et al. (2006) who developed a connectionist model including multiple levels of hierarchy. They assumed that ND can be judged based on representations on different levels of hierarchy and employed the model to explain the data on the effect of lesions of the perirhinal cortex on recognition memory. However, this model was not designed to have a high capacity for ND, and its capacity has not been tested.

A significant difference that distinguishes our approach from other computational models of ND is its generality. Instead of proposing a dedicated model for ND, we demonstrate in this work that existing predictive coding neural networks for AM or representation learning can perform robust (i.e., for images with correlated pixels), general (i.e., for sensory and semantic features) ND while maintaining a high capacity even with highly structured natural images.

### 4.2 Relationship to the predictive coding literature

In sections 3.2 and 3.3, we report results using rPCN to perform ND. The rPCN model we used (Tang et al., 2023b) is an ‘implicit’ variant of the original, ‘explicit’ formulation in Friston (2003, 2005), with naming convention coming from whether the variant represents the covariance matrix ‘implicitly’ or ‘explicitly.’ It is notable that the explicit variant theoretically should display slightly different behaviors on the same tasks, which we discuss in Appendix C.

The model we used in section 3.4 is slightly modified from Salvatori et al. (2021) by adding local connectivity constraints between layers 0 and 1. In their paper, Salvatori et al. (2021) demonstrated the ability of hPCN to perform a variety of AM tasks. However, the energy-based approach to ND we adopted in this paper is general and can potentially be applied to any energy-based model. In particular, a natural application of our approach is to temporal predictive coding network (tPCN), which is a multi-layer rPCN with a temporal dimension that has been shown to memorize videos/sequences of images (Tang et al., 2023a; Millidge et al., 2023). In this case, one way to detect novelty for sequences is to use, e.g., a running average of certain error neurons’ activities across time steps. Coincidentally, many studies have shown that there is a similar functional and anatomical overlap between ND for temporal order and other types of ND (see Warburton and Brown, 2015, for a review). Extending our current approach to tPCN can thus potentially fill this gap.

More generally, we predict the energy-based approach to generalize very naturally to any predictive coding models that have error neurons in their formulation. One flexible class of such models is proposed in Salvatori et al. (2022), where authors formulate predictive coding models on arbitrary graph topologies.

### 4.3 Relationship to experimental evidence

It has been suggested that the exact roles that the perirhinal cortex and hippocampus play in ND have partial overlaps (Brown and Aggleton, 2001). In particular, while lesion studies show that the perirhinal cortex plays a key role in ND for individual objects (Zola-Morgan et al., 1989; Meunier et al., 1993, 1996), hippocampal lesion only mildly impacts this ability (Honey et al., 1998). On the other hand, hippocampus damage has a greater effect when it comes to ND for the arrangement of individual objects (Gaffan and Parker, 1996) or novel pairing of individually familiar items (Aggleton and Brown, 1999) rather than individual objects themselves. The pattern is that, although different brain areas can be specialized in detecting one type of novelty, it also detects other types to some extent. This incomplete differentiation of functional roles is exactly what our results in Fig. 8C—here, even though layer 2 is highly specialized in detecting semantic novelty, it also detects (pixel-level) sensory novelty better than chance (i.e., a *d*^*′*^ separability of 0).

### 4.4 Relationship to anomaly & out-of-distribution detection

One key assumption we made in Fig. 6 for ND is that both familiar and novel patterns are samples from the same probability distribution. Data from the real world, however, likely comes from a multitude of probability distributions. When novel patterns can potentially be drawn from distinct, often unknown distributions, the task of distinguishing such novel patterns is known as out-of-distribution (OOD) detection or anomaly detection (AD) (Samariya and Thakkar, 2023; Ghamry et al., 2024) (for a survey and their exact relationship, see Yang et al. (2022)).

It is worth highlighting that, machine learning AD tasks/datasets such as MVTecAD (Bergmann et al., 2019) and various tasks involving CIFAR10 (Krizhevsky et al., 2012) necessarily require the detection of more subtle features beyond the simple, sensory (pixel-level) features that past ND computational models exclusively detect novelty for. Together with other desirable features such as local learning rules, PCNs have something to offer both as a putative model of brain circuits and as a machine intelligence algorithm to efficiently solve ND-related tasks.

### 4.5 Experimental predictions

Our PCN for ND model predicts that the neurons showing higher responses to novel stimuli should correspond to error neurons in PCN. Recently, more evidence for the existence of error neurons in early visual areas such as V2 (Huang et al., 2018) has emerged, and it has been observed that they have distinct genetic markers, paving way to new methods to identify them (O’Toole et al., 2023; Jordan and Keller, 2023). This opens up new experimental avenues to examine our prediction to identify novelty neurons as error neurons. It is noteworthy that this prediction is particularly robust to imprecise measurement because there is no need to consider fine-grained details at the level of individual neurons and how they encode information. All it needs is a sum/average activity across one layer afforded by current cell imaging (e.g., photometry).

Whether AM and ND are separable processes in the brain has been a consistent debate (Yonelinas, 2002; Yonelinas et al., 2010). Past literature has considered the functional difference between the hippocampus and perirhinal cortex as evidence to favor the dual-process theory (Brown and Aggleton, 2001). At least part of the difficulty causing the debate is the lack of clear definitions of *separable processes*. Without giving a clear answer, our model nevertheless provides some concrete grounding to think about the complex relationship—recognition memory can be thought of as a distributed process (across error neurons in different layers) in a circuit that performs ND on different levels of abstraction.

## Conclusions

This paper has proposed that ND can arise in PCNs, which provides an attractive hypothesis because **(1)** ND is performed by a general network that can also perform representation learning and AM; **(2)** it requires only local computations and plasticity (Tang et al., 2023b) and thus has a degree of biological plausibility; **(3)** it performs more robustly in ND tasks, especially when the patterns have an explicit correlated structure such as real-world images; and **(4)** our hierarchical model enables flexible ND for features of various abstraction levels. Moreover, we have shown analytically that this superior performance results from the covariance encoded in the recurrent weights of rPCN, stretching any query patterns and the mean of stored patterns accordingly before determining their novelty and familiarity. Overall, our work on ND combines recent advances in energy-based models for AM with experimental observation in neuroscience. This symbiosis eventually leads us to a biologically plausible, effective, and general computational mechanism underlying the discrimination of novel and familiar stimuli in the brain or artificial neural networks.

## Acknowledgments

This work has been supported by Medical Research Council UK grant MC UU 00003/1 to RB, an E.P. Abraham Scholarship in the Chemical, Biological/Life and Medical Sciences to MT. The authors would like to thank Gaspard Oliviers, Nima Mirkhani, Nicol S. Harper, Yuhang Song, Sumedha Nalluru and Mathilde Guillaumin for valuable feedback on the experiments and manuscript.

## Data availability

All code required for replicating the simulation presented in this paper can be found freely online at https://github.com/ltjed/novelty-detection-pc.

## Appendix

### A Proof of Theorem 1

To start, note that the training phase of rPCN can be seen as a constrained optimization problem: by Eq. 1, without loss of generality, assuming zero bias (***ν*** = 0), we have

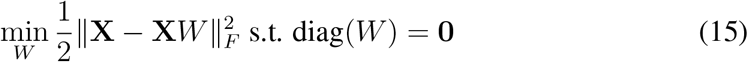

where for simplicity of notation, *W* in the appendix is the transpose of *W* used in the main text. Then, we can equivalently write the constraints into the Lagrangian

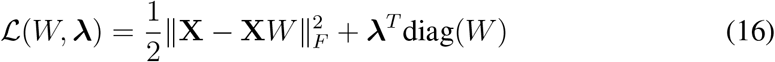

where ***λ*** = (*λ*_1_, …, *λ*_*d*_) is a vector of Lagrangian multipliers. Taking gradient with respect to *W* yields that,

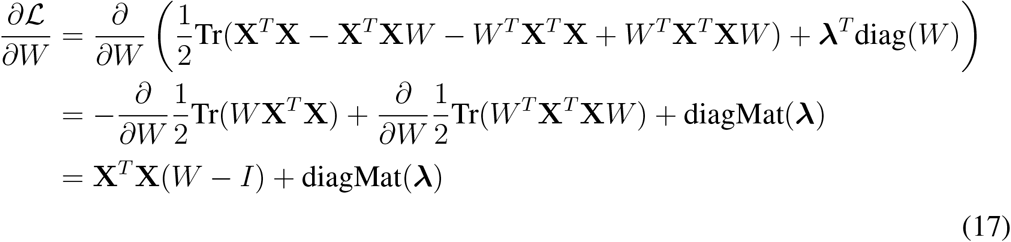

Similarly, taking gradient with respect to **λ** yields

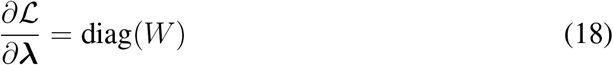

Setting the gradient 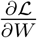 to **0** yields

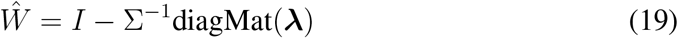

By substituting *Ŵ* into Eq. 18 and setting it to **0** we get

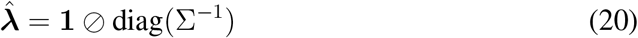

where ⊘ is the element-wise division. Finally, by substituting 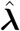 back into Eq. 19 we get the expression of the optimal *W* :

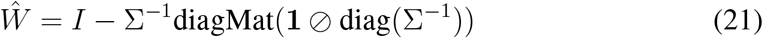

It can also be verified that (Ŵ, 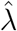) is indeed the global minimum by substituting it in Eq. 15.

Now, to express rPCN as performing metric learning in the form of Eq. 12, note that

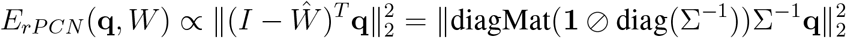

which concludes the proof.

#### B Details on the Experimental Procedure

To compare the model performances in Fig. 5, we

1. Draw *N* independent and identically distributed samples from the underlying data distribution as stored patterns as the training set for the model.
2. Draw *N* more independent and identically distributed samples from the underlying data distribution as novel patterns, each time making sure the samples are different from any of the *N* stored patterns through rejection sampling—rejecting until the sample drawn satisfies this requirement.
3. Feed a pair of patterns—one seen, one unseen—into the model as queries (i.e., keep the weights, *W*, constant) and evaluate each model’s energy value on these two patterns. A model’s judgment on this pair is correct if its energy value for the novel pattern is higher, and vice versa.
4. Repeat this step for all *N* seen-unseen pairs and calculate the error rate of a model as the number of incorrect judgments divided by *N*.

In particular, we calculate the *number of patterns retained N*_*retained*_ for the bottom row of Fig. 5 as

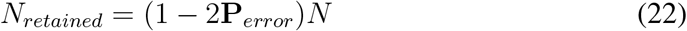

following Standing (1973), where **P**_*error*_ ∈ [0, 1] is the error rate.

#### C Explicit PCN Measures the Mahalanobis Distance

Explicit PCN (Friston, 2003) is another recurrent variant of PCNs that learns and encodes the covariance *explicitly* as parameters. Specifically, it encodes the subjective estimates of mean *µ*_true_ and covariance ∑_true_ with ***µ*** and **∑**, respectively. To improve its estimate, the model minimizes the *free energy*, which in this case is the negative multivariate Gaussian log-likelihood of the input pattern given the subjective parameters:

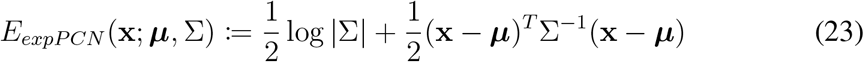

Like the derivations for HNs, we can ignore any terms that do not depend on the query **q**. Further, for simplicity, we also assume *µ*_true_ is **0** and that ***µ*** is a perfect estimate of it. This allows us to rewrite Eq. 23 as a function of **q** and **X**:

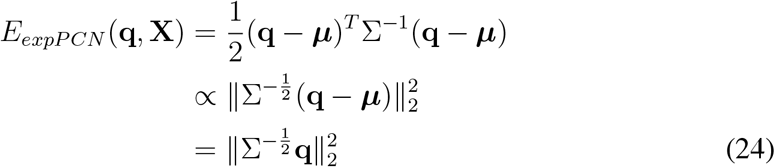

This is exactly the Mahalanobis distance, which is a well-known, *optimal* measure for distance in a correlated distribution Bellet et al. (2013), which effectively whitens the data and enables a fair comparison of (transformed) Euclidean distances.

Although the transformation performed by implicit PCN or rPCN (Fig. 9C) is not optimal when the query patterns are drawn from the same distribution that familiar patterns are sampled from, it can be more robust for out-of-distribution (OOD) detection. Consider the eigendecomposition of the covariance matrix, ∑ = *V* Λ*V* ^*T*^. For Fig. 9A, we have that *V* = (**v**_1_, **v**_2_), where **v**_1_ and **v**_2_ are unit vectors pointing towards the direction of familiar (purple) dot and novel (orange) point. The robustness of the implicit model to OOD detection can be seen by comparing the relative scaling effects along the principal components of the covariance matrix ∑—compared to exact whitening, implicit PCN is *less punishing* for variation along the first principal component, and *more punishing* for variation along the second (last) principal component. Since it follows from the Courant-Fischer theorem that samples from the distribution have the most variation along its first principal component and least variation along its last principal component, a sample **u** outside data distribution is likely to have larger 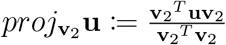 and thus classified as *more novel/surprising* by implicit PCN.

**Figure 9.**
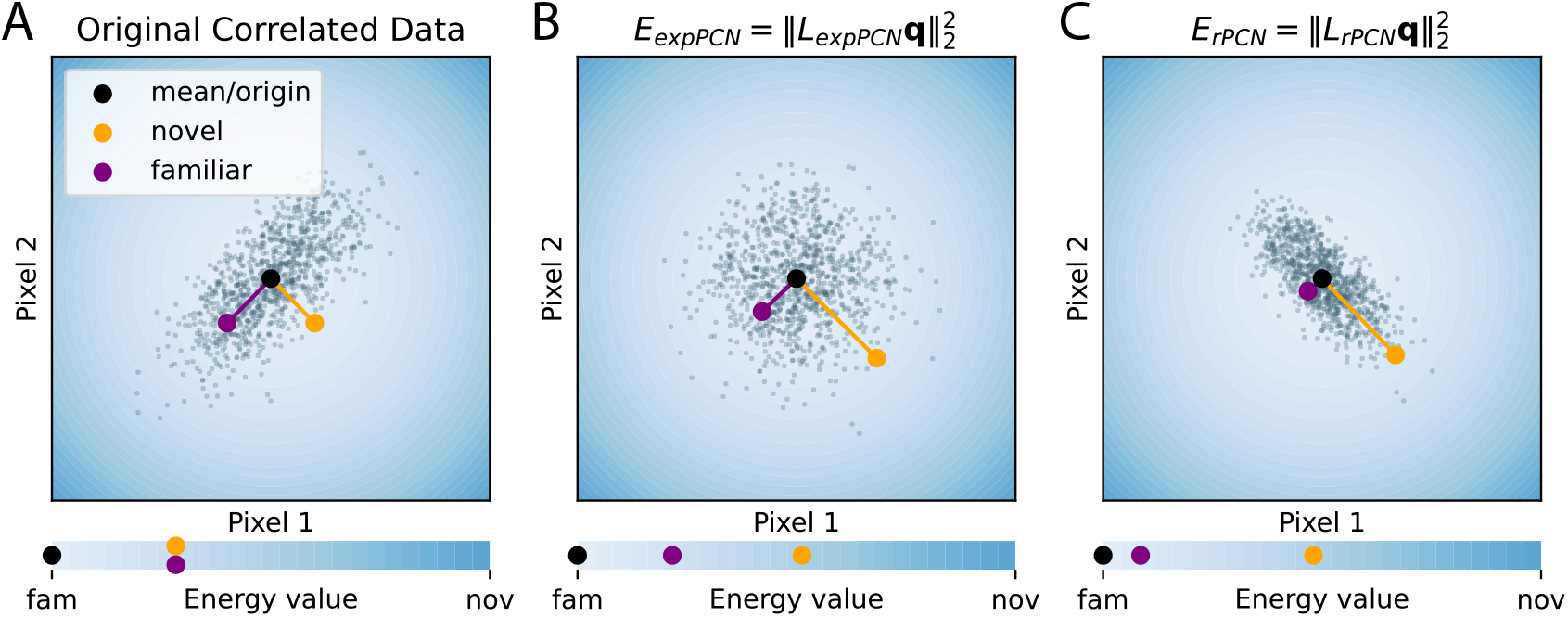
Comparing the effects of implicit and explicit PCN. Note that 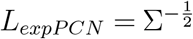 as derived in Eq. 24

